# Proliferation maintains the undifferentiated status of stem cells: the role of the planarian cell cycle regulator Cdh1

**DOI:** 10.1101/2021.06.13.448266

**Authors:** Yuki Sato, Yoshihiko Umesono, Yoshihito Kuroki, Kiyokazu Agata, Chikara Hashimoto

## Abstract

The coincidence of cell cycle arrest and differentiation has been described in a wide variety of stem cells and organisms for decades, but the causal relationship is still unclear due to the complicated regulation of the cell cycle. Here, we used the planarian *Dugesia japonica* since it possesses a quite simple cell cycle regulation in which *cdh1* is the only factor that arrests the cell cycle. When *cdh1* was functionally inhibited, the planarians could not maintain their tissue homeostasis and could not regenerate their missing body parts. While the ablation of *cdh1* caused pronounced propagation of the stem cells, the progenitor and differentiated cells were decreased. Further analysis indicated that the stem cells without *cdh1* expression did not undergo differentiation even though they received ERK signaling activation as an induction signal. These results suggested that stem cells could not acquire differentiation competence without cell cycle arrest. Thus, we propose that cell cycle regulation determines the differentiation competence and that cell cycle exit to G0 enables stem cells to undergo differentiation.

**Summary statement:** By using planarians, which have quite simple cell cycle regulation, we revealed that stem cells with cell cycle progression could not undergo differentiation even though an induction signal was activated.

## Introduction

Stem cells, including fertilized eggs, embryonic stem cells (ESCs), hematopoietic stem cells (HSCs), myoblasts and fibroblasts, have the ability of self-renewal and differentiation. Excessive differentiation causes the depletion of stem cells, and excessive proliferation causes carcinogenesis. Therefore, the precise balance of proliferation and differentiation is required for development, homeostatic tissue turnover and regeneration. It has been proposed that “induction” is primarily responsible for differentiation. For example, BMP-4 is required for epidermal differentiation of undifferentiated ectoderm (animal cap) cells in *Xenopus* early development (Sasai and De Robertis, 1997). The differentiation of sympathetic neurons from PC12 cells requires sustained ERK activation (Marhall, 1995; Avraham and Yarden, 2011). The activation of ERK signaling is also required for exit from pluripotency and for lineage commitment of mouse ESCs (Kunath et al., 2007; Stavridis et al., 2007). These reports emphasize the importance of induction signals to regulate the differentiation of stem cells. At the same time, the importance of “competence”, which is the capability to respond to induction signals has been proposed. For example, though the blastula animal cap cells of *Xenopus* have the ability to be induced by activin treatment to become mesoderm, the cap cells excised from the gastrula have lost this ability. Thus, both the extrinsic induction signals and the intrinsic competence are thought to be necessary for differentiation and they may function in parallel.

Meanwhile, it has been shown that the fate decision of stem cells such as myoblasts and embryonal carcinoma cells (ECCs) is made at the G1 phase (Nadal-Ginard, 1978; Wells, 1982). Notably, MyoD has already been present in proliferating myoblasts and bound to the regulatory region of its target genes, but the transcription is initiated only when the cell cycle is arrested (Skapek et al., 1995). Furthermore, human ESCs initiate the expression of marker genes for each germ layer at G1 phase, and the epigenetic status of these genes becomes bivalent at G1 (Pauklin and Vallier, 2013; Singh et al., 2015). These reports suggest that there is a mutual relationship between the cell cycle and differentiation, though the causal relationship remains to be clarified.

Cells decide whether to progress through or arrest their cell cycle during G1 phase. This decision depends on the level of cyclin-dependent kinase (CDK) activity. The CDK activity level becomes the lowest from late M phase to early G1 phase due to the degradation of Cyclins, and subsequent S phase entry requires the elevation of CDK activity. Especially, a ubiquitin E3-ligase, APC/C-Cdh1, is responsible for Cyclin degradation from the onset of G1 phase. To elevate the CDK activity for cell cycle progression, Cdh1 should be inactivated by binding of Emi1 or degraded through SCF-Skp2-mediated polyubiquitination. Therefore, the inactivation of APC/C-Cdh1 has been reported as a commitment point for progression through the cell cycle (Cappel et al., 2016). On the other hand, the sustained suppression of CDK activity induces cell cycle arrest at G1 phase. In addition to APC/C-Cdh1, CDK inhibitors (CKIs) such as INK family (p15, p16, p18 and p19) and CIP/KIP family (p21, p27 and p57) proteins inhibit CDK activity during G1 phase by direct binding. Furthermore, many other cell cycle regulators mutually interact with each other and construct a complicated network to regulate the cell cycle.

Previously, it was suggested that maintaining the proliferative state is important for the undifferentiated status of neural crest cells and lens placodal cells in *Xenopus* (Nagatomo and Hashimoto, 2007; Murato and Hashimoto, 2009), suggesting that cells tend to undergo differentiation once their cell cycle is arrested. If this is true, the cell cycle should be strictly regulated during development. It is known that cells can transiently arrest their cell cycle but can then proliferate again by restoring CDK activity during the developmental process. Such transient arrest makes cell cycle regulation highly dynamic. Since differentiation occurs at various times and places simultaneously during normal development in most multicellular organisms, complex and dynamic conversion of the cell cycle state seems to occur several times in the course of development. This is a major reason why investigation of the relationship between the cell cycle and differentiation has been problematic so far.

Planarians are well-known organisms that can regenerate any missing body part even from a tiny body fragment within a week. The remarkable regenerative ability is dependent on their adult stem cells called neoblasts (Agata and Watanabe, 1999; Newmark and Sánchez Alvarado, 2002). Since the neoblasts include pluripotent stem cells (PSCs), they give rise to all cell types in the planarian body, including germ-line cells (Shibata et al., 2010; Roberts-Galbraith and Newmark, 2015). By using the neoblasts, planarians also undergo perpetual tissue turnover throughout their life (Newmark and Sánchez Alvarado, 2000). The neoblasts are the only mitotic cells in the planarian body, and the cells never proliferate once they undergo differentiation. For this reason, planarian differentiation can be regarded as a kind of terminal differentiation. Since chemical inhibitors of ERK signaling and knockdown of the *erk-A* gene prohibit the differentiation of the neoblasts, the activation of ERK signaling is suggested to be responsible for the onset of differentiation as an induction signal (Tasaki et al., 2011a). Furthermore, planarians are one of the most basal organisms possessing three germ layers and three body axes, and are located at the branching point of protostomes and deuterostomes in the phylogenetic tree. Thus, universal features among multicellular organisms should have been acquired at planarians and conserved from them onwards.

Recently, genome analysis of multiple planarian species revealed that planarians have lost 124 genes essential for mice or humans (Grohme et al., 2018). *cdkn1b* encoding p27, a CIP/KIP family CKI, was one of the lost genes in planarians. It has also been reported that *cdkn1a* encoding p21 was not found in the genome of a European planarian, *Schmidtea mediterranea* (Pearson and Alvarado, 2010). Consistently, we could not find any CKI genes, including INK family and CIP/KIP family genes, in a draft genome of a freshwater planarian, *Dugesia japonica* (An et al., 2018). While most known factors regulating cell cycle arrest were not found in planarians, we found a *cdh1* gene (Fig. S1). Therefore, it could be thought that determination of whether cells arrest their cell cycle or enter the next round of cell division in planarians primarily depends on the presence or absence of Cdh1 expression. Interestingly, we also could not find Cdh1 inactivator genes *emi1* and *skp2* in the planarian genome. Taking these findings altogether, it is suggested that planarian cells cannot proliferate in the presence of Cdh1. Thus, it is possible to think that planarian species have a quite simple strategy for cell cycle regulation which is easy to manipulate by focusing only on *cdh1* expression. Furthermore, the neoblasts are the only proliferative cells in the planarian body, which enables us to focus on the cell cycle and differentiation of the neoblasts without any unintended effects on the other cells. Collectively, these considerations make planarians an ideal model for studying the relationship between the cell cycle and differentiation of stem cells.

In this study, we used *D. japonica* and showed that the ablation of planarian *cdh1* caused a drastic increase of the neoblasts, while the progenitor and differentiated cells were decreased. Moreover, the neoblasts without *cdh1* expression did not undergo differentiation, whereas ERK signaling was definitely activated in the course of regeneration. Based on these results, we propose a universal trait of stem cells that could be conserved among multicellular organisms: the cell cycle determines competence toward induction signals and only the stem cells arresting their cell cycle undergo differentiation according to the surroundings. This is the first report clearly showing that stem cells with cell cycle progression do not undergo differentiation even though they receive an induction signal.

## Results

### *cdh1* was expressed in the differentiating cells

While “induction” has attracted much attention, the importance of “competence” for differentiation has also been suggested for a long time. Accumulating evidence indicates that cell cycle arrest induces differentiation of stem cells (Lange and Calegari, 2010), but whether the cell cycle progression directly involves the repression of differentiation and maintenance of the undifferentiated state has been unclear. To investigate the relationship between the cell cycle and differentiation, we focused on the planarian *cdh1* gene, since it could be possible that *cdh1* plays a pivotal role in the decision of cell cycle arrest in planarians. Whole-mount in situ hybridization (WISH) revealed that *cdh1* was expressed in all body regions except the margin and pharynx (Fig. 1A). To examine whether the neoblasts expressed *cdh1*, we conducted double-WISH of *cdh1* and a neoblast marker gene, *piwiA*, (Yoshida-Kashikawa et al., 2007; Hayashi et al., 2010; Shibata et al., 2010). As a result, we observed that a fraction of the cells co-expressed *cdh1* and *piwiA* (Fig. 1B, white arrows) but we also observed single-positive cells of *cdh1* (Fig. 1B, black arrowheads) and *piwiA* (Fig. 1B, white arrowheads). Further, we examined the expression of *cdh1* during regeneration. WISH at 3 days post-amputation (dpa) revealed that *cdh1* was also expressed within the blastema region (Fig. 1C). Since the blastema region was formed by differentiating cells (*piwiA* mRNA negative/ PiwiA protein positive; Tasaki et al., 2011a), it is suggested that the *cdh1* was expressed in the committed neoblasts and differentiating cells. Based on these results and the known function of *cdh1*, planarian *cdh1* may be involved in cell cycle arrest in the differentiating cells.

**Fig. 1.**
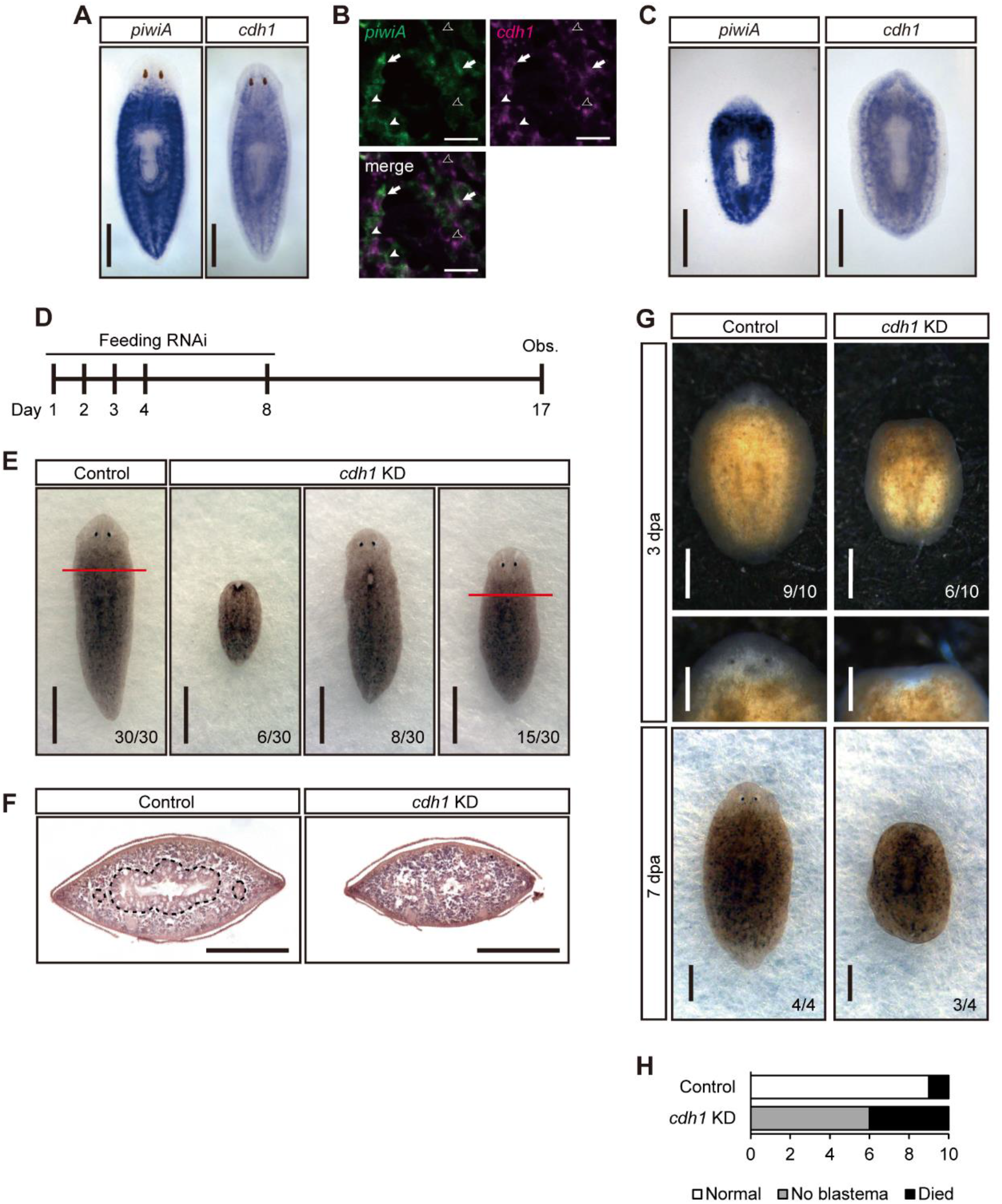
The expression pattern of *cdh1* and the phenotype of *cdh1* KD animals. (A) Expression pattern of *piwiA* and *cdh1* in intact planarian detected by whole-mount in situ hybridization. Cells expressing *cdh1* were observed at periphery of brain and mesenchymal space throughout the body. Scale bars, 1 mm. (B) Double fluorescent in situ hybridization of *piwiA* and *cdh1* in intact animals. Scale bars, 10 μm. White arrows indicate *piwiA/cdh1* double positive cells. White arrowheads indicate *piwiA* single positive cells. Black arrowheads indicate *cdh1* single positive cells. Scale bars, 25 μm. (C) Expression pattern of *piwiA* and *cdh1* at 3 dpa. *cdh1* expression was also detected in differentiating blastema region. Scale bars, 500 μm. (D) The schedule of feeding RNAi. The planarians were fed at day 1, 2, 3, 4 and 8. The phenotypes were observed at day 17. (E) The phenotypes of control and *cdh1* KD animals in tissue homeostasis. Whereas all control animals showed normal morphology, *cdh1* KD animals showed epithelium disorder or headless phenotype. Red lines indicate the section plane observed in Fig. 1F. Scale bars, 1 mm. (F) HE staining of transverse section anterior to the pharynx in control and *cdh1* KD animals. Dashed line indicates the morphology of intestine. The intestinal structure was collapsed in *cdh1* KD animals. Scale bars, 250 μm. (G) The phenotypes of control and *cdh1* KD animals undergoing regeneration at 3 dpa and 7 dpa. Control animals formed blastema at 3 dpa, and completely regenerated their lost tissues at 7 dpa. However, *cdh1* KD animals could not form blastema at 3 dpa, and failed to regenerate at 7 dpa. Scale bars, 500 μm in whole body samples. Scale bars, 250 μm in magnified view of regenerating head. (H) The number of individuals classified as each phenotype at 3 dpa.

### *cdh1* knockdown planarians showed disruption of homeostatic tissue turnover and blastema formation

To evaluate the function of *cdh1* in planarians, we conducted functional inhibition by feeding RNAi. We fed planarians dsRNA-containing food on 4 successive days from day 1 and once on day 8, and observed their phenotype at day 17 (Fig. 1D). Although the control animals maintained the normal body without any tissue disorder, some *cdh1* knockdown (KD) planarians had an epithelium disorder (8/30) or more severe headless phenotype (6/30), although the half of them (15/30) appeared normal (Fig. 1E). However, while the transverse section of the control animals showed normal morphology of the intestine, that of *cdh1* KD animals was collapsed even though the external morphology of the animals looked intact (Fig. 1F). Since planarians continuously undergo rapid tissue turnover, it is possible to think that the disruption of tissue homeostasis was caused by an insufficient supply of differentiated cells from the neoblasts.

We also examined the regeneration of *cdh1* KD animals after head and tail amputation at day 12 (Fig. 1G). The control animals formed blastemas normally and were regenerating their head and tail at 3 dpa. Notably, the head blastema of control animals was already regenerating eyes. On the other hand, more than half of *cdh1* KD animals (6/10) could not form blastemas, and the others had already died at 3 dpa (Fig. 1H). Finally, the control animals completely regenerated their missing body parts (4/4), but *cdh1* KD animals could not regenerate at all (3/4) at 7 dpa. Since blastema formation is the process that occurs subsequent to wound closure, we examined the wound healing response of *cdh1* KD animals by making an incision instead of amputation. In contrast to blastema formation, wound closure successfully occurred in 1 day both in control and in *cdh1* KD planarians, and JNK inhibitor SP600125 disturbed the wound closure, as previously described (Fig. S2; Tasaki et al., 2011b). This result indicated that the failure of blastema formation in *cdh1* KD animals was not a result of disruption of wound closure. Thus, it could be thought that the differentiating cells that form the blastema were not supplied in *cdh1* KD animals.

The disruption of tissue homeostasis and blastema formation both suggested that differentiated cells were not supplied sufficiently in *cdh1* KD animals. Such absence of differentiated cells could be explained by either the ablation of the neoblasts or the disruption of the differentiation process.

### The neoblasts were highly propagated in *cdh1* knockdown animals

To clarify whether the *cdh1* KD animals maintained their neoblasts, we checked the expression of neoblast marker genes. WISH of *piwiA* and *tgs1*, a candidate PSC marker gene in neoblasts (Zeng et al., 2018), showed that control and *cdh1* KD animals were indistinguishable at day 7, but the *cdh1* KD animals showed intense expression of both genes throughout their body at day 17 (Fig. 2A, B). This result can be explained in two ways: higher gene expression or increased number of the neoblasts. Then, we attempted to examine whether the neoblasts were increased by comparison of the areas where the neoblasts were present. Section in situ hybridization (SISH) at day 17 revealed that the area expressing *piwiA* within the total mesenchymal area was 18.3% in control animals and 46.1% in *cdh1* KD animals (Fig. 2C, D). Correspondingly, the percentage of mesenchymal area expressing *tgs1* was 1.98% in control animals and 16.3% in *cdh1* KD animals (Fig. 2C, D). The drastic increase of the area expressing neoblast marker genes indicated that the neoblasts were highly propagated in *cdh1* KD animals. Especially the area of PSCs (*tgs1*-positive cells) was more than 8.5 times larger.

**Fig. 2.**
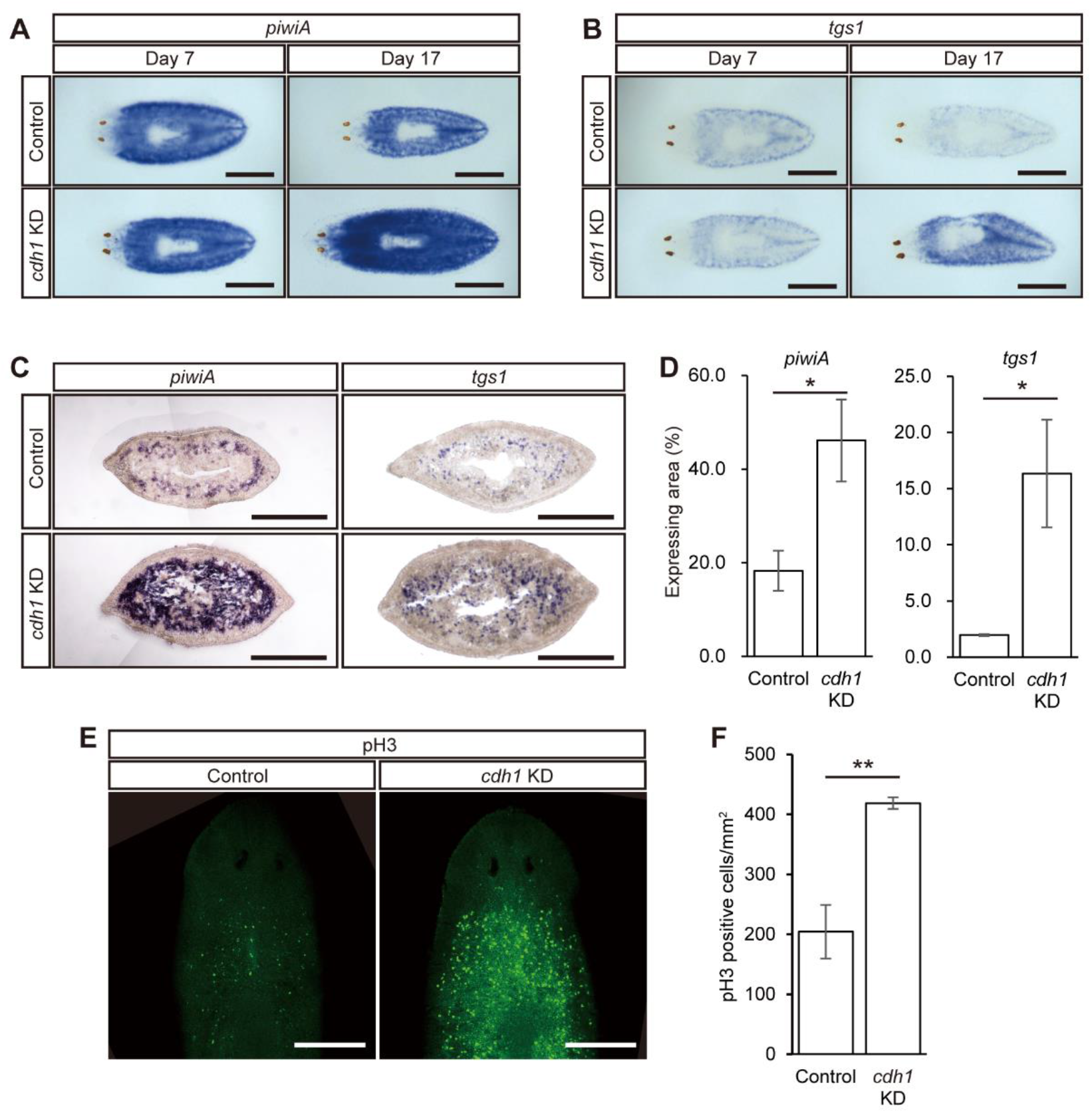
Functional inhibition of *cdh1* caused drastic increase of the neoblasts. (A) The expression pattern of *piwiA* in intact control animals and *cdh1* KD animals. *cdh1* KD animals showed more intense expression than control throughout the body at day 17. Scale bar, 500 μm. (B) The expression pattern of *tgs1* in intact control animals and *cdh1* KD animals. Like *piwiA, tgs1* was highly expressed in *cdh1* KD animals at day 17. Scale bar, 500 μm. (C) The expression pattern of *piwiA* and *tgs1* in transverse section anterior to the pharynx of control and *cdh1* KD animals at day 17. *piwiA*- and *tgs1*-expressing cells occupied a large area in *cdh1* KD animals. Scale bars, 250 μm. (D) Comparison of the *piwiA*- or *tgs1*-expressing area relative to total mesenchymal area in transverse section anterior to the pharynx of control and *cdh1* KD animals. Data are mean ± SEM (*n* = 3, Student’s *t*-test, \**p*<0.05). (E) Immunostaining of pH3 in control and *cdh1* KD planarians. *cdh1* KD animals showed an increased number of pH3-positive cells. Scale bars, 250 μm. (F) The number of pH3-positive cells in control and *cdh1* KD animals. Data are mean ± SEM (*n* = 3, Student’s *t*-test, \*\**p* < 0.005).

In addition to the increase of the area expressing neoblast marker genes, the number of phosphorylated histone H3 (pH3)-positive M phase cells was also increased in *cdh1* KD animals by more than 2-fold compared to that in control animals at day 17 (Fig. 2E, F), though the signal intensity looked much higher. Because the neoblasts are the only proliferative cells in the planarian body, the increase of pH3-positive cells also indicated an increase of the neoblasts in *cdh1* KD animals. These results suggested that the disruptions of both homeostatic turnover and regeneration in *cdh1* KD animals were caused not by the ablation of neoblasts but by the disruption of the differentiation process.

### The neoblasts in *cdh1* KD animals did not undergo differentiation in homeostatic condition

The drastic propagation of the neoblasts in *cdh1* KD animals suggested that the collapse of tissue homeostasis was caused by disruption of the differentiation process. To assess the differentiation of the neoblasts, we examined the expression of progenitor and differentiated cell marker genes by RT-qPCR analysis (Fig. 3A). Consistent with the results of in situ hybridization, the expression levels of *piwiA* and *tgs1* in *cdh1* KD animals were not distinguishable from those in control animals at day 7, but they started to be increased at day 11 and reached almost 3-fold higher than those in control animals at day 17. The S phase marker genes *pcna, mcm2* and *mcm3* also showed 3-4 fold higher expression at day 17 in *cdh1* KD animals than in control animals (Fig. 3A, Fig. S3). Taken together with the increase of M phase marker pH3-positive cells, this suggested the active cell cycling and accumulation of the neoblasts without any cell cycle arrest. In contrast to the neoblast marker genes, the expression of epithelial progenitor marker gene *prog1* and differentiated intestine marker gene *inx1* in *cdh1* KD animals was decreased to half or less of that in control animals (Fig. 3A). This suggested that the progenitor and differentiated cells were decreased in *cdh1* KD planarians. Since simple propagation of the neoblasts would also cause an increase of progenitor and differentiated cells, the decline of the progenitor and differentiated cells indicated that the process of differentiation was disrupted.

**Fig. 3.**
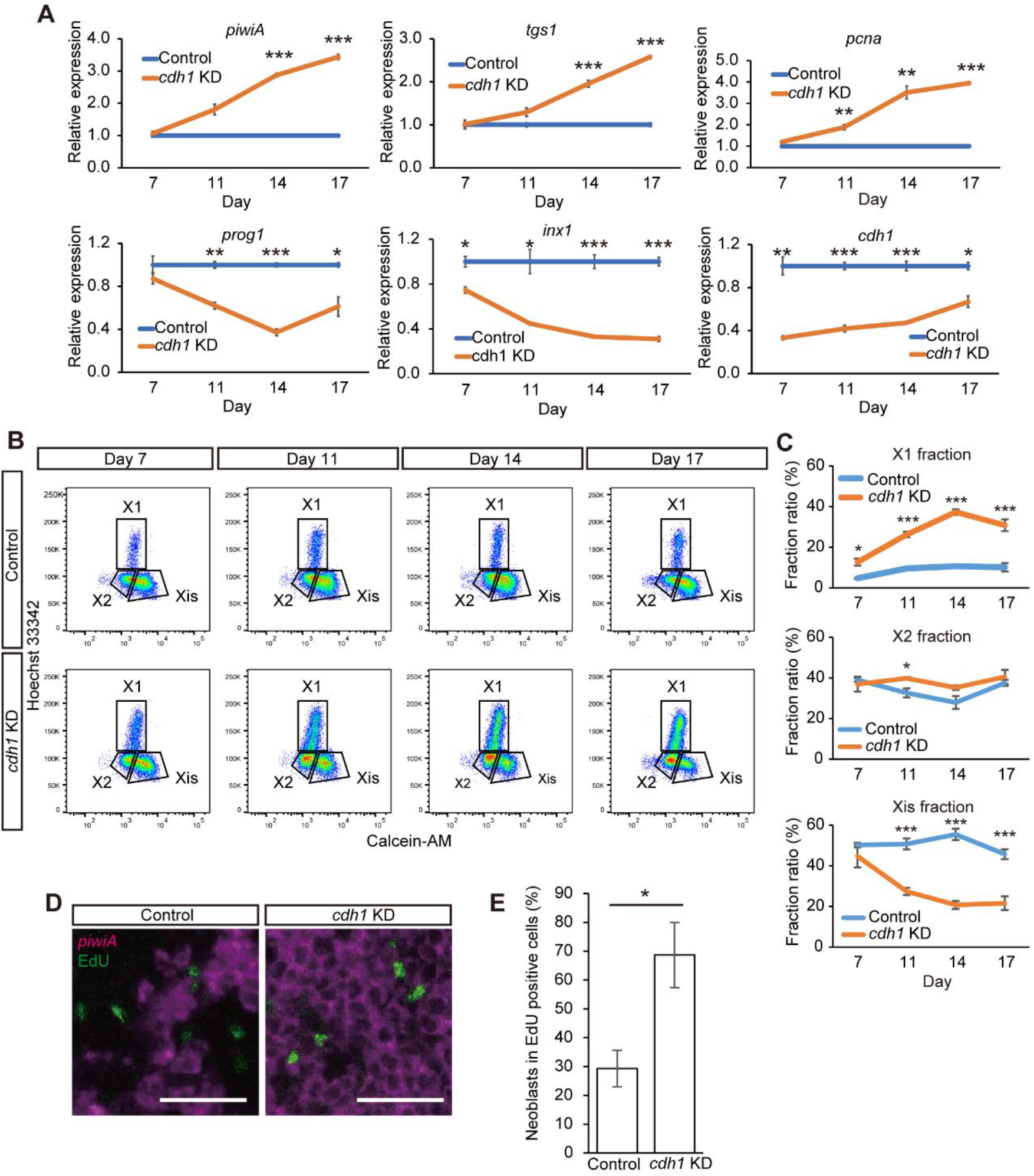
Functional inhibition of *cdh1* suppressed the differentiation of the neoblasts. (A) The gene expression levels in control and *cdh1* KD animals at day 7, 11, 14 and 17 determined by RT-qPCR analysis. The values were relative to in control animals. *cdh1* KD animals showed high expression of neoblast marker genes (*piwiA, tgs1* and *pcna*) but low expression of a progenitor gene (*prog1*) and a differentiated cell gene (*inx1*). Data are mean ± SEM (Student’s *t*-test, * *p* < 0.05, \*\**p* < 0.005, \*\*\**p* < 0.001). (B) Representative FACS profiles of cells derived from control and *cdh1* KD animals at day 7, 11, 14 and 17. The X-axis represents relative intensity of Calcein AM, which stains cytoplasm. The Y-axis represents relative intensity of Hoechst 33342, which stains nuclei. Cells that showed low intensity of Calsein AM and high intensity of Hoechst 33342 were designated the X1 fraction containing the neoblasts at S, G2 and M phase. The X2 fraction showing low intensity of Calcein AM and low intensity of Hoechst 33342 contains the neoblasts at G1 phase and a part of differentiated cells. Xis fraction showing high intensity of Calcein AM and low intensity of Hoechst 33342 contains only differentiated cells. *cdh1* KD animals showed an increase of X1 fraction and a decrease of Xis fraction. (C) Ratio of fraction of X1, X2 and Xis in control and *cdh1* KD animals. Data are mean ± SEM (Student’s *t*-test, * *p* < 0.05, \*\*\**p* < 0.001). (D) Fluorescent staining of *piwiA* and EdU at 3 days after EdU labeling in control and *cdh1* KD animals. Scale bar, 25 μm. (E) The percentage of *piwiA*-positive neoblasts in EdU-positive cells. More EdU-positive cells remained as the neoblasts in *cdh1* KD animals. Data are mean ± SEM (*n* = 3; Student’s *t*-test, * p < 0.05).

We also confirmed the change of cell populations by FACS (fluorescent-activated cell sorting) using nuclear and cytoplasmic staining by Hoechst33342 and Calcein AM (Hayashi et al., 2006), which distinguish the neoblasts from the differentiated cells independently of marker gene expression. FACS analysis classified the planarian cells into 3 fractions (Fig. 3B): the X1 fraction showing high nuclear content and scant cytoplasm contains the neoblasts in S, G2 and M phase, the X2 fraction showing low nuclear content and scant cytoplasm contains the neoblasts in G1 phase and a part of the differentiated cells, and the Xis fraction showing low nuclear content and developed cytoplasm contains only differentiated cells (Hayashi et al., 2006). The *cdh1* KD animals showed a higher ratio of the X1 fraction than the control animals from day 7 to 17 (Fig. 3B, C), which suggested that the enhancement of self-renewal by *cdh1* KD preceded the increase of neoblast marker gene expression at day 7. In contrast to the X1 fraction, the ratio of the Xis fraction became lower than that of the control at day 11, 14 and 17 (Fig. 3B, C). The total cell numbers of control and *cdh1* KD animals were comparable though the ratio between them was dynamically changed (Fig. S4), indicating that the neoblasts were propagated without differentiation while the differentiated cells were decreasing due to rapid tissue turnover. Of note, the density plot of *cdh1* KD animals showed a clear border between the X2 fraction and Xis fraction which could not be observed in that of control animals. It could be thought that the cells that were present between the X2 fraction and Xis fraction were differentiating progenitor cells. Therefore, the results also suggested the disappearance of differentiating progenitor cells while the neoblasts were propagated. The ratio of the X2 fraction was not so significantly changed from day 7 to day 17 (Fig. 3B, C). However, it could be thought that the differentiating cells disappeared while the neoblasts passing through G1 phase were increased within the X2 fraction, which made the X2 fraction seemingly unchanged.

For further understanding, we specifically labeled the neoblasts by EdU incorporation at day 12 and examined their differentiation at day 15 (Fig. 3D, E). In the control animals, 29.3% of EdU positive mesenchymal cells were *piwiA* positive neoblasts and the other 70.7% of EdU positive cells that had lost *piwiA* expression were newly differentiated cells. On the other hand, 68.7% of EdU positive mesenchymal cells retained *piwiA* expression in *cdh1* KD animals. This result clearly indicated that functional inhibition of *cdh1* suppressed the differentiation of the neoblasts. Taken together, these results indicated that the neoblasts were highly enriched but could not undergo differentiation in the homeostatic condition of *cdh1* KD planarians.

### *cdh1* was required for the neoblasts to respond to ERK signaling and undergo differentiation during regeneration

While control animals showed a differentiating *piwiA* negative blastema region at 3 dpa, *cdh1* KD animals did not have *piwiA* negative region (Fig. 4A). This regeneration defect resembled that of ERK inhibitor-treated animals (Tasaki et al., 2011a). ERK signaling has been thought to be an induction signal in planarians since an ERK inhibitor (U0126) and the knockdown of *erk-A* inhibited the differentiation of the neoblasts in both homeostatic and regenerating conditions (Tasaki et al., 2011a). Therefore, we speculated that the ERK signaling was not activated in the neoblasts in *cdh1* KD animals and caused the failure of blastema formation. However, surprisingly, western blotting analysis using antibody against the phosphorylated active form of ERK (pERK) revealed that pERK was detected near the stump of both control and *cdh1* KD animals at 1 dpa, while U0126 effectively inhibited the phosphorylation of ERK (Fig. 4B). To check the pERK activity in the neoblasts, we conducted double WISH of *piwiA* and *mkpA*, a reliable target gene of ERK signaling (Tasaki et al., 2011a; Umesono et al., 2013). *mkpA* was highly expressed near the stump at 1 dpa in control and *cdh1* KD animals, and U0126 treatment of control animals inhibited the *mkpA* expression (Fig. 4C). While most of the *mkpA* expression in control animals was observed in differentiated or differentiating *piwiA* negative cells, *mkpA* was largely expressed in *piwiA* positive neoblasts in *cdh1* KD animals. These results indicated that the ERK signaling was definitely activated in the neoblasts even in *cdh1* KD animals, but differentiation did not occur. To quantify the pERK activity, we conducted RT-qPCR analysis of *mkpA*. The result showed that the expression of *mkpA* was comparable between control and *cdh1* KD animals (Fig. 4D). Taken together, these results suggested that the neoblasts in *cdh1* KD animals could receive almost the same amount of ERK signaling activation, but they could not respond to the signal, and therefore the neoblasts could not undergo differentiation.

**Fig. 4.**
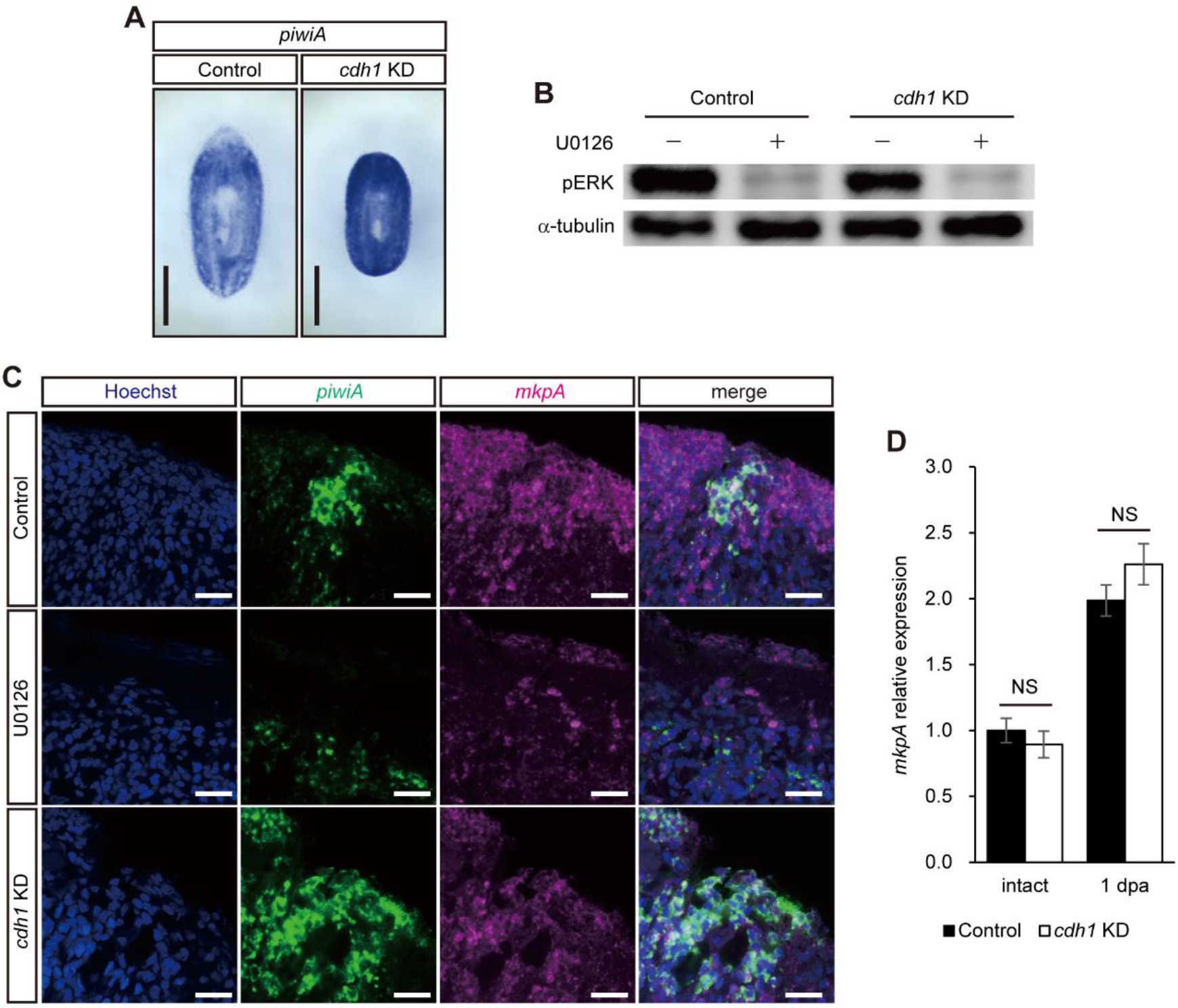
The neoblasts in *cdh1* KD animals did not form differentiating blastema but underwent activation of ERK signaling. (A) Expression pattern of *piwiA* in control and *cdh1* KD animals at 3 dpa. *cdh1* KD animals did not show differentiating blastema. Scale bars, 500 μm. (B) Western blotting of pERK in control and *cdh1* KD animals treated with DMSO or 25 μM U0126. *cdh1* KD animals showed pERK, like control animals, and U0126 effectively inhibited the phosphorylation of ERK in both animals. (C) Expression patterns of *mkpA* and *piwiA* in control and *cdh1* KD animals at 1 dpa. *mkpA* was expressed in differentiated or differentiating cells in control animals, and U0126 inhibited the expression. In contrast, the *mkpA* was expressed in the neoblasts in *cdh1* KD animals. Scale bars, 25 μm. (D) The expression level of *mkpA* in control and *cdh1* KD animals that were either intact or 1 dpa. The values are relative to those in control animals. The *mkpA* expression was comparable between control and *cdh1* KD animals. Data are mean ± SEM (Student’s *t*-test, NS: no significant difference).

Taken altogether, our findings show that the neoblasts in *cdh1* KD animals, which could not arrest the cell cycle, did not undergo differentiation even though they received an induction signal. Thus, our results indicate that cell cycle arrest is required by stem cells for the acquisition of competence to differentiate. Alternatively, it could be thought that stem cells persist in their undifferentiated state as long as their cell cycle is progressing.

## Discussion

In this report, we showed that planarians seem to lack most cell cycle regulator genes, but possess *cdh1* as their only gene causing cell cycle exit, suggesting that planarians could have a quite simple cell cycle regulation in which only Cdh1 expression regulated cell cycle exit. The *cdh1* KD planarians did not maintain tissue homeostasis and also did not regenerate their body after amputation. Under conditions that induced homeostasis in control planarians, the mitotic neoblasts in *cdh1* KD planarians, including PSCs, were drastically propagated while the progenitor cells and the differentiated cells were decreased. Furthermore, the neoblasts in *cdh1* KD animals could not undergo differentiation to form a blastema although the ERK signaling as an induction signal was definitely activated. These results indicated that *cdh1* KD enhanced self-renewal of the neoblasts, and the neoblasts in the proliferative state could not undergo differentiation.

Our results summarized above strongly indicated that proliferative stem cells could not respond to induction signals. In other words, it is possible to think that maintenance of the proliferative state ensures the undifferentiated state. Interestingly, Tgs1, a candidate marker for PSCs among neoblasts (Zeng et al., 2018), carries APC/C-Cdh1 target sequences such as a D-box. This indicates that Tgs1 would be degraded, and then cells would lose the identity of PSCs upon the expression of *cdh1*, which supports a direct association between pluripotency and cell cycling. Further, animal cap cells of *Xenopus* blastulae are pluripotent, but the pluripotency disappears at the onset of gastrulation, excluding the prospective neural crest region. This is consistent with the previous report that neural crest cells retain blastula-stage potential during *Xenopus* early development (Buitrago-Delgado et al., 2015). Coincidently, though *Xhairy2* is weakly expressed in the entire animal cap region in blastulae, the *Xhairy2* expression is restricted only in the prospective neural crest region and becomes higher at gastrula stage or later (Tsuji et al., 2003). Since *Xhairy2* represses *p27* expression to maintain cells in a proliferative and undifferentiated state, continuous expression of *Xhairy2* from the blastula through the gastrula stage maintains the cell cycling, and this may ensure the pluripotency (Nagatomo and Hashimoto, 2007).

The incompatibility of proliferation and differentiation has been described in a wide variety of stem cells and organisms for a long time. In *C. elegans* gonad, germ-line stem cells (GSCs) exist only adjacent to the distal tip cell (DTC) and undergo differentiation when they are apart from DTC. It is known that the self-renewal of GSCs in *C. elegans* is maintained by the activation of Notch signaling by DTC, and that all GSCs undergo differentiation in recessive mutants of GLP-1, a Notch homolog in *C. elegans* (Austin and Kimble, 1987). Similarly, GSCs in male *Drosophila melanogaster* exist only when they contact hub cells through E-cadherin. The GSCs that contact hub cells receive Unpaired and activate JAK-STAT signaling to proliferate, while GSCs not in contact with hub cells cannot receive a sufficient amount of Unpaired and they therefore undergo differentiation (Kiger et al., 2001; Tulina and Matunis, 2001). Although different molecular pathways are activated in the GSCs of the above two invertebrate species, the GSCs share the feature that they exist as stem cells in a microenvironment promoting self-renewal, and undergo differentiation when self-renewal cannot be maintained. This raises the possibility that the difference of the activated signaling pathway in stem cells is due to the difference of how to maintain the stem cells’ cell cycle progression. Consistently, *cyclin D* KD planarians, which are thought to undergo forced cell cycle arrest, could not regenerate their missing body parts and lost the neoblasts (Fig. S5). These reports and our results indicate that the cell cycle should be maintained for stem cells to exist in tissues without differentiation.

The incompatibility of proliferation and differentiation is also reported in the regulation of transcription factors and chromatin status. Cyclin D-CDK and cyclin E-CDK complexes phosphorylate Oct4, Sox2 and Nanog to stabilize these transcription factors from proteasomal degradation in mouse ESCs, suggesting that cell cycle progression maintains a stem cell state (Liu et al., 2017). Conversely, the transcriptional activity of MyoD, a master regulator of muscle differentiation whose ectopic expression causes the expression of the muscle-specific genes in many cell types, is activated only at G1 phase in murine myoblasts (Skapek et al., 1995). The proneuronal differentiation factor NGN-2 is phosphorylated by Cyclin A- and Cyclin B-CDK complexes and this phosphorylation impairs the DNA-binding of NGN2 during S through M phase in mice and *Xenopus* (Ali et al., 2011; Hindley et al., 2012). Moreover, H3K4me3 active histone marks are increased at G1 phase, while H3K27me3 repressive histone marks are stable throughout the cell cycle at developmentally regulated genes of human ESCs. This suggested that the developmentally regulated genes become bivalent at only G1 phase, which allows the transcription factors to be loaded (Singh et al., 2015). Our results also showed that the neoblasts in *cdh1* KD planarians could not undergo differentiation though they had phosphorylated ERK (Fig. 4). These examples indicate that stem cells in the proliferative state could not undergo differentiation even if they expressed potent differentiation-specific transcription factors.

In addition, stalk cell differentiation of cellular slime mold *Dictyostelium* requires cell cycle arrest. This suggests that the incompatibility between proliferation and differentiation is a universal trait of cells beyond the animal kingdom. Interestingly, DIF-1, an inducer of stalk cell differentiation in *Dictyostelium*, suppresses the expression of *cyclin D* and *cyclin E* and induces the differentiation of mouse vascular smooth muscle cells and the re-differentiation of mouse leukemia cells (Miwa et al., 2000; Asahi et al., 1995). Furthermore, we observed upregulation of *mkpA* in planarians treated with DIF-1. It could be thought that DIF-1 also induced the activation of ERK as induction signal in planarians (Fig. S6A). However, DIF-1 treatment of *cdh1* KD planarians did not cause blastema formation, which confirmed that stem cells with cell cycle progression could not undergo differentiation even though the induction signal was artificially activated (Fig. S6B).

Taking these facts all together, we propose that stem cells may persist in their undifferentiated state as long as their cell cycle is progressing, and cell cycle arrest might dictate that stem cells become competent to differentiate according to their surroundings. Although the activated signaling pathways and transcription factors vary among stem cells, it could be thought that the incompatibility of proliferation and differentiation is a universal feature conserved from *Dictyostelium* to mammals.

It has been reported that differentiation occurs at G1 phase in various stem cells and that differentiation is induced by functional inhibition of Cyclins or CDKs (Lange and Calegari, 2010). These reports make us speculate that stem cells stochastically undergo differentiation depending on whether their cell cycle phase is G1 or not. The differentiation at G1 phase could possibly explain the stochastic differentiation model which has been proposed for ESCs and a range of adult tissue homeostasis (Stumpf et al., 2017, Simons and Clevers, 2011). If this speculation is correct, the proliferating neoblasts in *cdh1* KD animals also stochastically undergo differentiation during G1 phase in the ongoing cell cycle. However, the differentiated cells and the progenitor cells were definitely decreased in *cdh1* KD animals, which caused disruption of tissue homeostasis and regeneration. Since *cdh1* is known to play an important role in cell cycle arrest, it could be suggested that differentiation occurs when the cell cycle is arrested but not during the progressing G1 phase.

As mentioned above, *Xhairy2* maintains the proliferative and undifferentiated state by repressing *p27* during neural crest specification and also in the lens placode of *Xenopus* embryo (Nagatomo and Hashimoto, 2007; Murato and Hashimoto, 2009). However, the prospective neural crest region or lens placode did not show frequent cell division compared to the other regions of the embryo. Thus, it could be thought that these cells do not completely stop the cell cycle but just pause at G1 phase. Reversible growth arrest has been described just as “quiescence” so far, but here we would like to propose two substantially different quiescent states, namely, G1 arrest and G0: cells in G1 arrest might be just transiently paused but still remain in G1 phase to persist in their undifferentiated state, but cells in G0 phase might completely stop proliferation and exit the cell cycle for the acquisition of competence. Although we could not distinguish between G1 arrest and G0 yet, we expect that further research on the cell cycle and differentiation competence will delineate the difference underlying the two quiescent states.

During the development of most multicellular organisms, stepwise differentiation occurs in parallel at various time points and regions. According to our model described above, cells must exit the cell cycle in order to undergo differentiation, and re-enter the cell cycle to increase the cell number. This combination of steps of proliferation and differentiation may correspond to the seeming stepwise differentiation. Furthermore, multiple mechanisms regulating cell cycle exit should be required for parallel and spatiotemporal differentiation. Such a requirement for complicated cell cycle regulation could possibly explain why most multicellular organisms conserve numerous factors involved in cell cycle regulation. However, it was suggested that planarians have a quite simple cell cycle regulation which lacks most of the cell cycle regulators such as CKIs. The differentiated cells in planarians are supplied directly from the neoblasts and never proliferate, which might allow the simplification of cell cycle regulation. In addition, cell cycle regulation without CKI expression is also observed in mouse ESCs. Due to the absence of CKI expression, CDK activity is constitutively high in mouse ESCs, and their G1 phase is quite short (about 3 hours) compared to that of mouse embryonic fibroblasts (11 hours; Liu et al., 2019). Therefore, it is conceivable that planarians could have lost CKI genes which are dispensable for cell cycle regulation of PSCs. In other words, planarian cell cycle regulation might not be species-specifically anomalous, but instead might be regulation common among PSCs. Studies of planarians, which have simplified cell cycle regulation, will reveal the nature of features of proliferation and differentiation conserved among multicellular organisms.

The remarkable regenerative ability of planarians has fascinated many researchers, and the availability of methods for functional inhibition by RNAi in planarians provides a great opportunity to study gene function in planarians. However, the lack of a means for gain-of-function in planarians still makes these studies difficult. For this purpose, effective isolation and amplification of PSCs is needed. In this report, we showed the drastic enrichment of the neoblasts throughout the body of *cdh1* KD animals, which could be regarded as “culture flasks of the neoblasts”. Therefore, the functional inhibition of *cdh1* may enable us to establish not only the gain-of-function but also gene knockout, conditional mutagenesis and other genetic manipulations, which could be a key to the next step of planarian research.

## Acknowledgements

We thank Dr. Kazuyuki Fujimitsu for helpful discussions, Miyuki Ishida for technical assistance and critical discussions, and Dr. Elizabeth Nakajima for critical reading of the manuscript.

## Author contributions

Conceptualization: Y.U., C.H.; Methodology: Y.S., Y.K.; Validation: Y.S.; Formal analysis: Y.S., Y.K.; Investigation: Y.S.; Resources: K.A., C.H.; Writing-original draft preparation: Y.S., C.H.; Supervision: C.H.; Project administration: Y.U., C.H.; Funding acquisition: C.H.

## Materials and methods

### Animals

The clonal SSP-9T strain of the planarian *Dugesia japonica* (Nishimura et al., 2015), derived from the Iruma River in Gifu prefecture, Japan, was maintained in dechlorinated tap water at 21°C, which is a suitable condition for maintaining the population size (Mori et al, 2019). Chicken liver was fed to them one or two times a week. Planarians ~6 mm in length were starved for at least 1 week before experiments. The animal experimentation was conducted according to the protocol reviewed and approved by the Institutional Animal Care and Use Committee of JT Biohistory Research Hall.

### Molecular cloning of planarian *cdh1* gene

The transcriptome dataset of *D. japonica* (Shibata et al., 2016) was used for gene identification, and we thereby found the planarian *cdh1* gene (accession number: IAAB01050803). cDNA of the gene was obtained by PCR using a set of primers (Fw: 5’-ATGGATAGTTCATATGAACGTCGATTATT-3’ and Rv: 5’-TTATCTCATACCACTGAACAAATCGAGAGC-3’), and cloned into pCS2 vector and sequenced.

### Inhibitor treatment

The JNK inhibitor SP600125 (Sigma-Aldrich) and the MAPK/ERK (MEK) inhibitor U0126 (Cell Signaling Technology) were dissolved in dimethylsulfoxide (DMSO) as 25 mM. DIF-1 (Sigma-Aldrich) was dissolved in DMSO at 500 μM. Planarians were kept in dechlorinated tap water containing 0.1% DMSO, 25 μM SP600125, 25 μM U0126 or 500 nM DIF-1 with light shielding. The inhibitor-containing breeding water was replaced with fresh inhibitor-containing breeding water every day.

### Whole-mount in situ hybridization

Planarians were treated with 2% hydrochloric acid in 5/8 Holtfreter’s solution for 5 minutes at room temperature to remove mucus and fixed with 4% paraformaldehyde, 5% methanol in 11/14 PBS for 30 minutes at room temperature. Hybridization and staining of digoxigenin (DIG)- or fluorescein (FITC)-labelled probes were conducted as described previously (Umesono et al., 1997). After the hybridization, samples were washed and treated with 1% blocking reagent (Roche). For alkaline phosphatase staining, samples were incubated with anti-DIG-AP antibody (Roche, 11093274910) overnight, and color development was conducted with BCIP/NBT substrate (Roche). For fluorescent staining, anti-DIG-POD (Roche, 11207733910) or anti-FITC-POD (Roche, 11772465001) antibody was used for overnight incubation. To optimize the fluorescent staining, we washed samples with borate buffer (100 mM borate (pH 8.5), 0.1% Tween-20), and conducted tyramide signal amplification in TSA reaction solution (a tyramide reagent, 0.003% H_2_O_2_, 2% dextran sulfate sodium salt and 0.3 mg/mL 4-iodo-phenol in borate buffer) according to Lauter et al., 2011 and Akiyama-Oda and Oda, 2016.

### Whole-mount immunostaining of pH3

Samples were fixed and immunostaining was conducted according to a previous report (Tasaki et al., 2011a). Anti-pH3 antibody (Sigma-Aldrich, 06-570) was used at 1/200 dilution as first antibody, and anti-rabbit IgG-Alexa 488 (Invitrogen, A11034) was used at 1/1,000 dilution as secondary antibody.

### Section in situ hybridization

Planarians were treated with 2% hydrochloric acid in 5/8 Holtfreter’s solution for 5 minutes at room temperature to remove mucus and fixed with relaxant solution (1% nitric acid/ 1.6% formaldehyde/0.02 mM MgSO_4_) overnight at 4°C. The following procedures were conducted as previously described (Kobayashi et al., 1998).

### Feeding RNAi

Double-stranded RNA (dsRNA) was synthesized based on a previous report (Rouhana et al., 2013). A DNA fragment containing the target gene sequence with the T7 promoter at both ends was used as the template for dsRNA synthesis. The synthesis was conducted with a Megascript T7 transcription kit (Thermo Fisher Scientific) following the manufacturer ‘s protocol. The synthesized dsRNA was treated with TURBO DNase and RNase T1 (Roche) for 1 h at 37°C to remove DNA template and single-stranded RNA. The dsRNA was precipitated with LiCl and dissolved in water at 250 ng/μl. Thirty planarians were fed a mixture of 25 μL of 50% chicken liver homogenate (w/v), 5 μL of 2% agarose (w/v), and 10 μL of dsRNA solution, once daily for 3 h. The mixture was divided into small aliquots and frozen at −30°C before feeding. The feeding was conducted on four successive days and on the day of 1 week after the first feeding. Control animals were fed dsRNA of *enhanced green fluorescent protein* (*EGFP*).

### RT-qPCR analysis

Total RNA was extracted from 3-5 planarians by using ISOGEN-LS (Wako) following the manufacturer’s protocol. cDNA was synthesized from 1 μg total RNA using a PrimeScript RT reagent kit (Takara) according to the manufacturer’s instructions. The synthesized cDNA was diluted 1:20, and qPCR reactions were conducted in 20 μL of a mixture containing 1x TB Green premix Ex TaqII (Takara), 0.3 μM gene-specific forward/reverse primers and 2 μL of diluted cDNA using a thermal cycler Dice (Takara). The reactions were carried out as follows: 95°C for 30 sec, 40 cycles of 95°C for 15 s, 60°C for 30 s, 72°C for 1 min. The expression of each gene was normalized by the expression of *g3pdh*. All experiments were performed using three biological and three technical replicates. The sequences of gene-specific primers are listed in table S1.

### FACS analysis

Cell dissociation was performed from ten planarians as previously described (Hayashi et al., 2006). The number of dissociated cells was counted using a LUNA-FL (Logos Biosystems). The dissociated cells were stained with 18 μg/mL Hoechst 33342 and 0.1 μg/mL Calcein AM for 2 h at 20°C. FACS analysis was conducted using blue and violet lasers of BD FACSMelody (BD Bioscience) and FlowJo (v10.7.1, BD Bioscience) (Kuroki, unpublished). Experiments were performed using three biological replicates.

### EdU labeling and detection

Thirty planarians were fed a mixture of 20 μL of 50% chicken liver homogenate (w/v), 5 μL of 2% agarose (w/v), 10 μL of the indicated dsRNA solution and 14 μL of 20 mg/mL EdU for 3 h. The mixture had been divided into small aliquots and frozen at −30°C before feeding. The animals were fixed and subjected to whole-mount in situ hybridization as described above. After that, EdU detection was conducted using a Click-iT EdU imaging kit (Invitrogen) according to the manufacturer’s protocol.

### Western blotting

Blastemas were dissected from 20 regenerating planarians at 1 dpa, pooled, and dissolved in 50 μl of sample buffer (0.01 M Tris-HCl, 2% SDS, 6% 2-mercaptethanol, 10% glycerol) and were sonicated and boiled for 5 minutes. 5 μl of the samples was subjected to SDS-PAGE and western blotting. Membrane blocking was conducted by using Blocking Reagent (Roche). The primary antibodies were rabbit anti-phosphorylated ERK (1/500; Tasaki et al., 2011a) or mouse anti-α-tubulin monoclonal antibody DM 1A (1/5,000; Sigma-Aldrich, T9026). An appropriate antibody conjugated with horseradish peroxidase (1/5,000) was used as secondary antibody. ImmunoStar Zeta (Wako) was used for signal detection.

### Phylogenetic analysis

The amino acid sequences of Cdh1 from various organisms and CDC20 from *Drosophila melanogaster* were aligned using the ClustalW program. A phylogenic tree was reconstructed from the alignment by the neighbor joining method with the JTT model included in Mega X software.

### Statistical analysis

Statistical analyses were performed using Microsoft Excel. Two-sided Student’s *t*-tests (α = 0.05) were performed to compare the means of two populations.

